# Structural insight into the individual variability architecture of the functional brain connectome

**DOI:** 10.1101/2022.02.16.480803

**Authors:** Lianglong Sun, Xinyuan Liang, Dingna Duan, Jin Liu, Yuhan Chen, Xindi Wang, Xuhong Liao, Mingrui Xia, Tengda Zhao, Yong He

## Abstract

Human cognition and behaviors depend upon the brain’s functional connectomes, which vary remarkably across individuals. However, whether and how the functional connectome individual variability architecture is structurally constrained remains largely unknown. Using tractography- and morphometry-based network models, we observed the spatial convergence of structural and functional connectome individual variability, with higher variability in heteromodal association regions and lower variability in primary regions. We demonstrated that functional variability is significantly predicted by a unifying structural variability pattern and that this prediction follows a primary-to-heteromodal hierarchical axis, with higher accuracy in primary regions and lower accuracy in heteromodal regions. We further decomposed group-level connectome variability patterns into individual unique contributions and uncovered the structural-functional correspondence that is associated with individual cognitive traits. These results advance our understanding of the structural basis of individual functional variability and suggest the importance of integrating multimodal connectome signatures for individual differences in cognition and behaviors.

## 1. Introduction

Each individual thinks or behaves differently from other individuals. A growing number of studies have suggested that the brain’s functional connectomes (FCs) and structural connectomes (SCs) act as biological substrates underlying these individual differences in cognition and behaviors (Bullmore and Sporns, 2009; Liao et al., 2017; Park and Friston, 2013). Several prior studies have documented that the intrinsic FC profiles of the human brain during resting conditions vary across individuals, with higher variability primarily in heteromodal association regions and lower variability in primary sensorimotor and visual regions (Mueller et al., 2013; Gao et al., 2014; Xu et al., 2018; Stoecklein et al., 2019). Such an FC variability pattern is significantly correlated with focal cortical folding variation (Mueller et al., 2013; Mansour et al., 2021) and evolutionary cortical expansion (Mueller et al., 2013; Stoecklein et al., 2019) and predicts individual differences in cognitive domains (Mueller et al., 2013; Liao et al., 2017). In the present study, we investigated whether and how interindividual FC variability is anatomically shaped by the SC variability of the human brain.

There are currently two principal methods available for mapping the SCs of the human brain: white matter tractography-based methods using diffusion weighted imaging and gray matter morphometry-based methods using structural imaging. Specifically, tractography-based connectomes can be reconstructed by inferring axonal tracts among brain regions using deterministic or probabilistic tractography approaches (Gong et al., 2009; Hagmann et al., 2008; Zhang et al., 2022), while morphometry-based connectomes can be obtained by examining the statistical similarity of morphometric measures among regions (He et al., 2007; Tijms et al., 2012; Kong et al., 2015). Notably, the morphometry-based connectomes do not directly measure physical pathways between regions but provide complementary information for the measurement of axonal projections (Gong et al., 2012; Evans, 2013; Seidlitz et al., 2018). Although there are mounting reports showing SC-FC coupling across brain regions or subjects (Honey et al., 2009; Baum et al., 2019; Wang et al., 2015b; Misic et al., 2016; Zimmerman et al., 2018), whether and how interindividual FC variability patterns are structurally constrained is understudied. To date, only three studies have examined the relationship between interindividual FC and SC variability patterns. Of them, two studies reported nonsignificant spatial correlations between FC variability and tractography-based SC variability (Chamberland et al., 2017; Karahan et al., 2021), which could be attributable to a relatively small sample size (n = 9 (Chamberland et al., 2017) and n = 29 (Karahan et al., 2021)) and confounds of intraindividual variation (Karahan et al., 2021). A very recent study reported a significant correlation between interindividual variability patterns of FC and tractography-based SC (Mansour et al., 2021). However, there are several important issues that have not been addressed. First, these previous studies used only a single SC feature in which network edges represent the existence of direct white matter connections, ignoring the contribution of network communications (such as path length (Achard et al., 2006) and communicability (Crofts and Higham, 2009) and morphometry-based SCs, which are important anatomical scaffoldings for shaping the brain’s functional activities (Goni et al., 2014; Avena-Koenigsberger et al., 2017; Alexander-Bloch et al., 2013; Geng et al., 2017). Second, these prior studies focus mainly on the relationship between whole-brain SC and FC variability, ignoring the spatial heterogeneity of brain regions. Third, considering that there are structural and functional variabilities in brain networks even within an individual, thus the estimation and control of intrasubject variability are important to accurately depict the relationship between SC and FC variability at both group and individual levels. Thus, the correspondence between interindividual FC and SC variability has been difficult to establish to date.

To fill these gaps, we conducted a comprehensive integrative analysis of multimodal connectome features to unravel the relation between functional and structural variability in the human brain using repeated-measures functional, structural, and diffusion imaging data from the Human Connectome Project (HCP) (Van Essen et al., 2013). Specifically, we first calculated the canonical interindividual FC variability in the whole brain and then quantified multifaceted interindividual SC variability patterns that captured three communication models of tractography-based connectomes and five morphometric measures of morphometry-based connectomes. Next, we used both linear and nonlinear computational analyses to test the hypothesis that individual FC variability is structurally constrained by unifying SC signatures across the whole brain and across different systems. Finally, we decomposed the group-level FC and SC variability patterns into individual unique contributions and then explored the alignment of structural-functional uniqueness and its relevance to individual cognitive and behavioral traits.

## 2. Materials and Methods

### 2.1. Participants and Data Acquisition

#### Participants

We used two publicly available multimodal magnetic resonance imaging (MRI) datasets from the Human Connectome Project (HCP) (Van Essen et al., 2013). For detailed subject inclusion/exclusion criteria, please refer to (Van Essen et al., 2013). The HCP S1200 dataset included 1012 healthy young-adult subjects (ages 22-37, 543 females) with complete minimal preprocessed imaging data for all modalities. The HCP Test-Retest (TRT) dataset included 42 subjects (ages 22-35, 30 females) who underwent two separate scans with an interval ranging from 0.5 to 11 months. The HCP TRT dataset was used as the discovery dataset. The HCP S1200 dataset was used as the validation dataset. Written informed consent was obtained from all subjects, and the scanning protocol was approved by the Institutional Review Board of Washington University in St. Louis, MO, USA (IRB #20120436).

#### Data Acquisition

All MRI data were acquired on a customized 3 T 32-channel Siemens Skyra scanner at Washington University. All images in the HCP S1200 dataset and HCP TRT dataset shared the same scanning parameters. Resting-state functional MRI (rs-fMRI) images were obtained by multiband gradient-echo-planar imaging acquisitions with two rs-fMRI runs (the phase encoding direction corresponded to left-to-right and right-to-left, respectively). The sequence parameters for each run were the same as follows: repetition time (TR) = 720 ms, echo time (TE) = 33.1 ms, flip angle = 52°, bandwidth = 2290 Hz/pixel, field of view = 208 × 180 mm^2^, matrix = 104 × 90; 72 slices, voxel size = 2 × 2 × 2 mm^3^, multiband factor = 8, and 1200 volumes. The high spatial resolution diffusion-weighted imaging (DWI) data (1.25 mm isotropic, 18 b0 acquisitions, 270 diffusion-encoding directions with three shells of b=1000, 2000, and 3000 s/mm^2^, 90 directions for each shell, 2 × 2 × 2 mm isotropic voxels, TR = 5520 ms, TE = 9.58 ms) were acquired by using a Stejskal-Tanner diffusion-encoding scheme. T1-weighted (T1w) images were acquired using a 3D-magnetization-prepared rapid acquisition with gradient echo (MPRAGE) sequence (0.7 mm isotropic voxels, matrix = 320 × 320; TR = 2400 ms, TE = 2.14 ms, 256 slices, flip angle = 8°).

T2-weighted (T2w) images were acquired using a 3D T2-sampling perfection with application-optimized contrasts by using flip angle evolution (SPACE) sequence with identical geometry (TR = 3200 ms, TE = 565 ms).

### 2.2. Data Preprocessing

#### 2.2.1. Functional data

All functional imaging data were preprocessed by the HCP minimal preprocessing pipeline (Glasser et al., 2013), including gradient distortion correction, motion correction, echo-planar imaging distortion correction, registration to the Montreal Neurological Institute (MNI) space, and intensity normalization. Then, the volume time series were mapped to the standard CIFTI grayordinates space, downsampled to 32k_fs_LR mesh, and slightly smoothed using a 2 mm full-width half-maximum (FWHM) kernel on the surface. As a part of the preprocessing pipeline, ICA-FIX denoising was used to remove nonneural spatiotemporal noise and head motion. To further reduce the effects of nuisance covariates, we regressed out the white matter, cerebrospinal fluid, global signals, and the 12 head motion parameters and performed temporal bandpass filtering (0.01-0.1 Hz) using SPM12 (https://www.fil.ion.ucl.ac.uk/spm/) and GRETNA (Wang et al., 2015a).

#### 2.2.2. Diffusion data

All diffusion imaging data were preprocessed with the HCP diffusion preprocessing pipeline, including mean b0 image normalization, echo planar imaging (EPI) distortion correction, eddy-current distortion correction, head motion correction, gradient nonlinearity correction, linear registration to native structural space using a 6 degrees of freedom (DOF) boundary-based registration, and data masking with the final brain mask to reduce the file size (Glasser et al., 2013).

#### 2.2.3. Morphological data

All T1w and T2w MRI scans went through the HCP structural preprocessing pipeline (Glasser et al., 2013). We obtained the individual cortical thickness, cortical curvature, sulcal depth, surface area, and intracortical myelination after bias correction in the standard surface (32k_fs_LR space) from the publicly available dataset. Intracortical myelination is characterized by the ratio of the T1w value to the T2w value (Glasser and Van Essen, 2011).

### 2.3. Network Reconstruction

#### 2.3.1. Functional connectome (FC)

A surface-based multimodal brain atlas (HCP-MMP1.0) was used to parcellate the cerebral cortex into 180 regions of interest (ROIs) per hemisphere (Glasser et al., 2016). For each run, we first obtained the mean time series of vertices in each brain node and calculated Pearson’s correlation coefficient of the time series between any pair of nodes to construct the FC. Then, we performed Fisher’s r-to-z transformation to normalize the correlation coefficient of the individual FCs.

#### 2.3.2. Tractography-based structural connectome (SC)

We transformed the surface-based parcel labels into each individual’s native volume space by using HCP Workbench’s command *label-to-volume-mapping*. These atlas labels at volume space were further dilated by 2.5 mm to enter the gray matter-white matter boundary. Together, 360 dilated regions represented the nodes of SCs.

The edges of the direct white matter SCs were defined using probabilistic tractography. Briefly, the preprocessed b0 data and crossing fiber modeled diffusion data (called BedpostX data) were obtained from the HCP database. Each defined brain node was selected as a seed region, and probabilistic tractography was performed by sampling 5000 streamline fibers for each voxel within each seed region. The connectivity probability from the source region to the target region was defined by the number of streamlines passing through the target region divided by the total number of streamlines sampled from the source region. One challenge of probabilistic tractography is that the number of streamlines drops with distance from the seed mask, which may underestimate long-distance connections in the whole-brain network. Thus, we performed the distance correction using the --pd flag in fsl ProbtrackX tools. As a result, the connectivity weight is the expected length of the pathways times the streamlines number (Behrens et al., 2007; Cui et al., 2013). The tractography procedure was repeated for all pairs of brain regions to obtain a whole-brain weighted connectivity matrix. After performing the symmetrization operation, the tractography-based direct SC was generated for each individual. The above procedures were implemented with the FSL (Jenkinson et al., 2012) and the PANDA Toolkit (Cui et al., 2013).

The tractography-based direct SC represents the direct communication between brain regions, while convergent evidence has emphasized that signal propagation among brain regions may also occur along one or more indirect pathways (Crofts and Higham, 2009; Goni et al., 2014; Suárez et al., 2020; Vazquez-Rodriguez et al., 2019). To characterize interindividual structural variability from the perspective of different regional communication models, we derived another two weighted connectomes that characterized two types of multipath communication mechanisms. As two extremes of the polysynaptic communication models among brain regions, path length (PL) characterizes the routing protocols of information propagation, and communicability (CO) reflects the diffusion processes of information propagation.

##### Path length

The cost of a connection in the weighted SC network was first calculated by inverting the edge weight, i.e. *Cost_ij_* = 1/*A_ij_*, where *A_ij_* is the weight of the edge between node *i* and *j*. The path length is defined as the minimum cost of the contiguous edges between two nodes.

##### Communicability

Communicability is defined as the weighted sum of all walks between two nodes (Crofts and Higham, 2009; Estrada and Hatano, 2008). For a binary network A, communicability *CO_ij_* is defined as

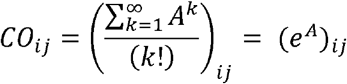

where (*A^k^*)_*ij*_ represents the number of walks within k steps that start at node *i* and finish at node *j*.

Note that the walks of step *k* are normalized by a penalty factor 1 /(/*k*!) to ensure that the shorter the walk is, the greater the contribution. For a weighted network *A*, the communicability *C_ij_* is defined as

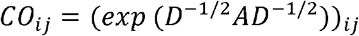

where *D* is a diagonal degree matrix formed by *D* ≔ *diag*(*d_i_*), and *d_i_* is the generalized degree of node *i* formed by 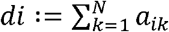. Path-length-based SC and communicability-based SC were implemented using the GRETNA toolbox (Wang et al., 2015b) and netneurotools (https://netneurotools.readthedocs.io), respectively.

#### 2.3.3. Morphometry-based SC

We use a common framework (Kong et al., 2015; Li et al., 2021b; Tijms et al., 2012) to map individual morphometry-based SCs based on the interregional similarity of morphological features. Considering that the distribution of morphological features of brain regions is discrete, we choose the earth mover’s distance, which is a statistical measurement of the difference between two discrete probability distributions (Elizaveta Levina, 2001). Specifically, for each morphological feature, including cortical thickness, cortical curvature, sulcal depth, surface area, and intracortical myelination, we calculated the earth mover’s distance between the feature distributions of any pair of regions and then obtained a dissimilarity matrix. We normalized each dissimilarity matrix to the range [0,1] and quantified the interregional similarity as 1-distance. Finally, we obtained five morphological-based SCs for each individual, with large values in those SCs indicating high morphological closeness.

### 2.4. Estimating Interindividual Variability Patterns at the Group Level

Following the method proposed by Mueller and colleagues (Mueller et al., 2013), we calculated the adjusted interindividual variability in brain connectomes. Taking the FC as an example, the raw interindividual FC variability of a given brain region *i* was defined as follows:

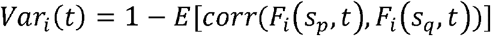

where *p,q* = 1,2 … *N*(*p* ≠ *q*); *N* is the number of subjects in the dataset; *s_p_* and *s_q_* indicate the subject; and *t* indicates the separated session. *F_i_*(*s_p_, t*) is the functional connectivity profile of region *i* of subject s_*p*_ in session *t*. Similarly, when calculating the individual SC variability, *F_i_* is the structural connectivity profile between region *i* and all other brain regions.

For each subject, the intraindividual variance in region *i* was estimated using repeat-scan data from all sessions/runs,

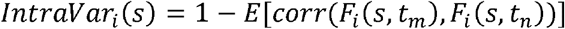

where *m,n* = 1, … *T; T* is the total number of sessions/runs. Then, by averaging the intraindividual variance of *N* individuals, we obtained the intraindividual variability of region *i* that can characterize the entire dataset,

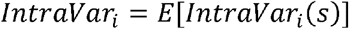

To estimate the adjusted interindividual variability, we regressed out the intraindividual variability map from the raw interindividual variability map using a general linear model and then averaged the interindividual variability map across sessions/runs.

### 2.5. Principal Component Analysis

Considering distinct structural patterns provided by tractography-based SCs and morphometry-based SCs, we first generated a linear multivariate representation of the structural variability by performing a principal component analysis (PCA) of the interindividual variability maps of all eight structural features. Specifically, the SC variability maps of all features were first z-transformed and combined into a region × feature matrix *X*. Then, PCA was applied to *X*, yielding principal axes and the coefficients of linear combinations via the eigenvalue decomposition of the covariance matrix. The principal components were defined as the coordinate representations of features on the corresponding principal axes. The first principal component of SC variability captured the largest proportion of total variances. We calculated the ratio of the squared factor score (Abdi and Williams, 2010) of each feature by the eigenvalue associated with the principal component to characterize its contribution to the principal component.

### 2.6. Predicting Interindividual FC Variability Using a Multivariate Prediction Model

We employed a support vector regression (SVR) model (*fitrsvm* function in MATLAB) with a ‘*rbf*’ nonlinear kernel to examine the ability of SC variability to predict FC variability. Specifically, the FC variability map for the whole brain was represented as a vector (*N_regions_* × 1). All eight SC variability maps were chosen as input features (*N_regions_* × *M_metrics_*)· We first linearly scaled all features to the range of 0-1. Then, the prediction model was trained and evaluated using the leave-one-out cross-validation (LOOCV) strategy. The variability map of each region across the whole cortex was designated as the testing sample, while the variability maps of the remaining regions were defined as the training samples. After LOOCV, Pearson’s correlation coefficient between the predicted FC variability and the observed FC variability across all regions was calculated as the prediction accuracy. The same predictive framework was also performed for regions within each hierarchical system separately.

### 2.7. Direct SC-FC Coupling

Inspired by Bertha and colleagues (Vazquez-Rodriguez et al., 2019), we performed a multiple regression linear model to fit the FC profile in each region using SC profile predictors of all structural metrics in the same region. The regional SC-FC coupling was quantified by the fitted adjusted *R^2^*, which represents the degree of correspondence of structural and functional connectivity profiles. For each FC and SC metric, a group-level network was first estimated as the averaged connectivity matrix across individuals and sessions. Then, the estimated FC profile of node *i* was represented as

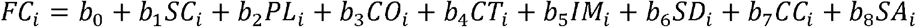

where the independent variables *SC_i_,PL_i_*, and *CO_i_* are the tractography-based structural connectivity between node *i* and all other nodes, and *CT_i_, IM_i_, SD_i_, CC_i_*, and *SA_i_* are the morphometry-based structural connectivity between node *i* and all other nodes. Such a multilinear regression framework to characterize SC-FC correspondence has been widely reported (Betzel et al., 2019; Goni et al., 2014; Vazquez-Rodriguez et al., 2019).

### 2.8. Mediation Analysis

To investigate whether the effect of SC variability on FC variability was mediated by the regional distribution of SC-FC coupling, a bootstrapped mediation analysis was employed using the MATLAB package *Mediation ToolBox* (https://github.com/canlab/MediationToolbox). We first normalized the independent (*X*, principal SC variability), dependent (*Y*, functional variability), and mediating (*M*, SC-FC coupling) variables. Then, we examined the total effect of principal SC variability on FC variability (path *c*), the relationship between SC-FC coupling and principal SC variability (path *a*), the relationship between FC variability and SC-FC coupling (path *b*), and the direct effect of principal SC variability on FC variability controlling for the mediator (i.e., SC-FC coupling) (path *c*’) The significance of the mediation/indirect effect (*ab*) of principal SC variability on FC variability through the mediator was tested using a bootstrapping analysis (resampled 10,000 times). For the SVR model-based result, we repeated the mediation analysis using predicted FC variability instead of principal SC variability.

### 2.9. Spatial Permutation Testing (Spin Test)

To further test whether spatial distributions of FC and SC variabilities follow a hierarchical manner from primary to heteromodal organization, we used a spherical projection null framework to estimate whether these variabilities are determined by the hierarchical classes or derived by spatial autocorrelation (Alexander-Bloch et al., 2018; Liu et al., 2020). Specifically, we first stratified all 360 cortical regions into four cortical functional hierarchies (Liu et al., 2020) so that each vertex in the cortical surface had a hierarchy type assignment. Mean variability values were then calculated within each hierarchical class. Under the premise of preserving spatial autocorrelation, the class labels were randomly rotated in the spherical space of the cortical surface, and the mean variability values were recomputed. After 10,000 permutations, the class-specific mean variability values were expressed as z scores relative to this null model. A positive z score indicated greater variability than expected by chance, and a negative z score indicated smaller variability than expected by chance. The p value of each class was defined as the proportion by the mean variability values in the null model that exceeded the true mean variability value. We also used this spin test to assess the significance of the alignment between two brain spatial patterns at both group and individual levels, including the correspondence between observed FC and SC variability pattern, the observed FC and predicted FC variability pattern, and the predicted FC and SC variability pattern.

### 2.10. Estimating Individual Uniqueness at the Subject Level

#### 2.10.1. Individual deviation quantification for each subject

For a given subject *s*, we estimated the deviation of individual connectivity profiles from the population-level connectivity profiles for each FC and SC by decomposing the definition of individual variability at the group level (Mueller et al., 2013). Specifically, we first computed the raw interindividual deviation of brain region *i* by estimating the dissimilarity of the connectivity profile of subject *s* with the profile of all other subjects as follows in the equation below. This measurement gives an overall description of the degree of uniqueness between the connectivity profile of the current node and that of other nodes.

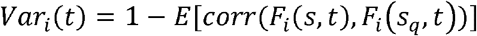

where *q* = 1,2, …, *N; N* is the number of subjects in the dataset; *s_q_* indicates the subject; and *t* indicates the session. *F_i_*(*s_q_, t*) is the connectivity profile of region *i* of subject *s_q_* in session *t*. After calculating the raw interindividual deviation of each region, we obtained an interindividual deviation map for each subject. Second, we calculated the intraindividual variance map *IntraVar*(*s*) for subject *s*. Finally, we regressed out the intraindividual variance from the raw interindividual deviation map to estimate the real individual deviation map of subject *s*. This subject-specific connectivity deviation could act as an indicator to characterize individual brain uniqueness. Together, we obtained the individual uniqueness of one FC and eight SCs in each session/run for this given subject.

For each subject, the individual-level correspondence between SC uniqueness and FC uniqueness was estimated following the same analysis procedure as previously described at the group level.

#### 2.10.2. Within-subject reliability of individual uniqueness

To investigate whether individual FC and SC uniqueness were stable within subjects across repeated sessions and variable between subjects, we performed the following reliability analysis and individual identification analysis.

##### Reliability analysis

For a given subject, we evaluated the within-subject similarity by calculating Pearson’s correlation coefficient of individual uniqueness in two sessions (S1 to S2). Next, we quantified the between-subject similarity by averaging the Pearson’s correlation of the uniqueness pattern between this given subject in S1 (S2) and all other subjects in S2 (S1). All subjects then had within- and between-subject similarity metrics. The individual uniqueness is considered to be reproducible if the within-subject similarity is significantly larger than the between-subject similarity. We performed a nonparametric permutation test to explore whether the reproducibility of individual uniqueness was significantly larger than random levels. Briefly, the subjects were randomly permutated, and the difference between within- and between-subject similarity was recomputed. This permutated procedure was repeated 10,000 times, yielding a null distribution. The p value was defined as the proportion of permutations with a difference value that exceeded the value in the observed data.

##### Individual identification analysis

We implemented the individual identification procedure as described by Finn and colleagues (Finn et al., 2015). Briefly, for all individual deviation maps in both sessions (S1 and S2), if the deviation maps from two repeated sessions of a given individual showed the highest similarity among any other two pairs, the identification was correct. The success ratio was calculated as the fraction of individuals who were identified correctly. The individual identification analysis was performed in two directions, one from S1 to S2 and the other from S2 to S1. Finally, we averaged the success ratio across these two directions.

### 2.11. Relevance for Cognition and Behavioral Performance

#### 2.11.1. Cognitive and behavioral measurements

We obtained the independent behavioral phenotypes in the HCP S1200 dataset provided by Tian and colleagues (Tian et al., 2020). In brief, 109 raw behavioral and cognitive measurements in the HCP dataset were selected for each subject, involving alertness, cognition, emotion, motor, personality, sensory, psychiatric and life function, substance use, and in-scanner task (emotion task, gambling task, language task, relational task, social task, and work memory task). Subjects missing one or more measurements were excluded from the analysis (final sample size n = 958). Applying a data-driven independent component decomposition pipeline, the 109 behavioral and cognitive measures were parsed into five independent summarizing dimensions, including cognition, illicit substance use, tobacco use, personality and emotional traits, and mental health (detailed in Tian et al., 2020). These five summarizing measurements were used in the following brain-behavior analysis.

#### 2.11.2. Partial least-squares (PLS) analysis

We then performed a partial least-squares correlation analysis with the myPLS toolbox (https://github.com/danizoeller/myPLS) to evaluate the implications of the structural-functional uniqueness correspondence in individual cognitive and behavioral performance. As a data-driven multivariate statistical technique, PLSC analysis was widely used to delineate the brain-behavior association by performing singular value decomposition (SVD) to obtain the orthogonal latent components (LCs) (Krishnan et al., 2011). LCs are the optimal linear combinations of the original variables from two matrices that maximize their covariance. Specifically, for brain domains, we considered the structural-functional uniqueness alignments at both the whole-brain and system levels. For behavioral domains, we used the five behavioral measurements mentioned above. We regressed out age and sex from brain measurements and behavioral measurements. Then, after z-scoring brain data *X* (subjects × brain measures) and behavioral data *Y* (subjects × behavioral measures) across all subjects, we computed the brain-behavior covariance matrix *R*,

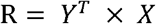

followed by performing SVD on *R*,

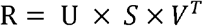

where *U* and *V* are the singular vectors, which could be called brain and behavioral weights, and S is a diagonal matrix containing the singular values. Therefore, each LC includes a distinct brain weight and a distinct behavioral weight. By linearly mapping the original brain and behavioral measurements of each subject onto their respective weights, the subject-specific brain and behavioral composite scores were estimated. To determine the significance level of each LC, we conducted a permutation test as follows. By performing 10,000 permutations to the brain measurements (randomly reordering the subjects) and leaving behavioral measurements unchanged, we calculated 10,000 null brain-behavior covariance matrices and obtained a sampling distribution of the singular values under the null hypothesis. The statistical significance of each LC was computed by comparing the singular value of the observed LC with its null distribution. For LC interpretation, we obtained the brain (behavioral) loadings by calculating Pearson’s correlation coefficients between the original brain (behavioral) measurements and brain (behavioral) composite scores. A large positive (or negative) loading for a given brain (behavioral) measurement indicates greater importance of this brain (behavioral) measurement for the LC. Using bootstrap resampling (1000 iterations), we computed 95% confidence intervals for the brain loadings and behavioral loadings.

## 3. Results

### 3.1. FC and SC Individual Variability Patterns Represent Cortical Hierarchical Organization

We leveraged multimodal functional, structural, and diffusion imaging data from 42 subjects with repeated scans in the HCP Test-Retest (TRT) dataset (Van Essen et al., 2013). For each individual, we reconstructed the FCs and eight types SCs of the brain (Fig 1A). For FC and each type of SC, we calculated intra- and inter-individual variability patterns according to a classic approach proposed by Mueller and colleagues (Mueller et al., 2013) (Fig 1B).

**Figure 1.**
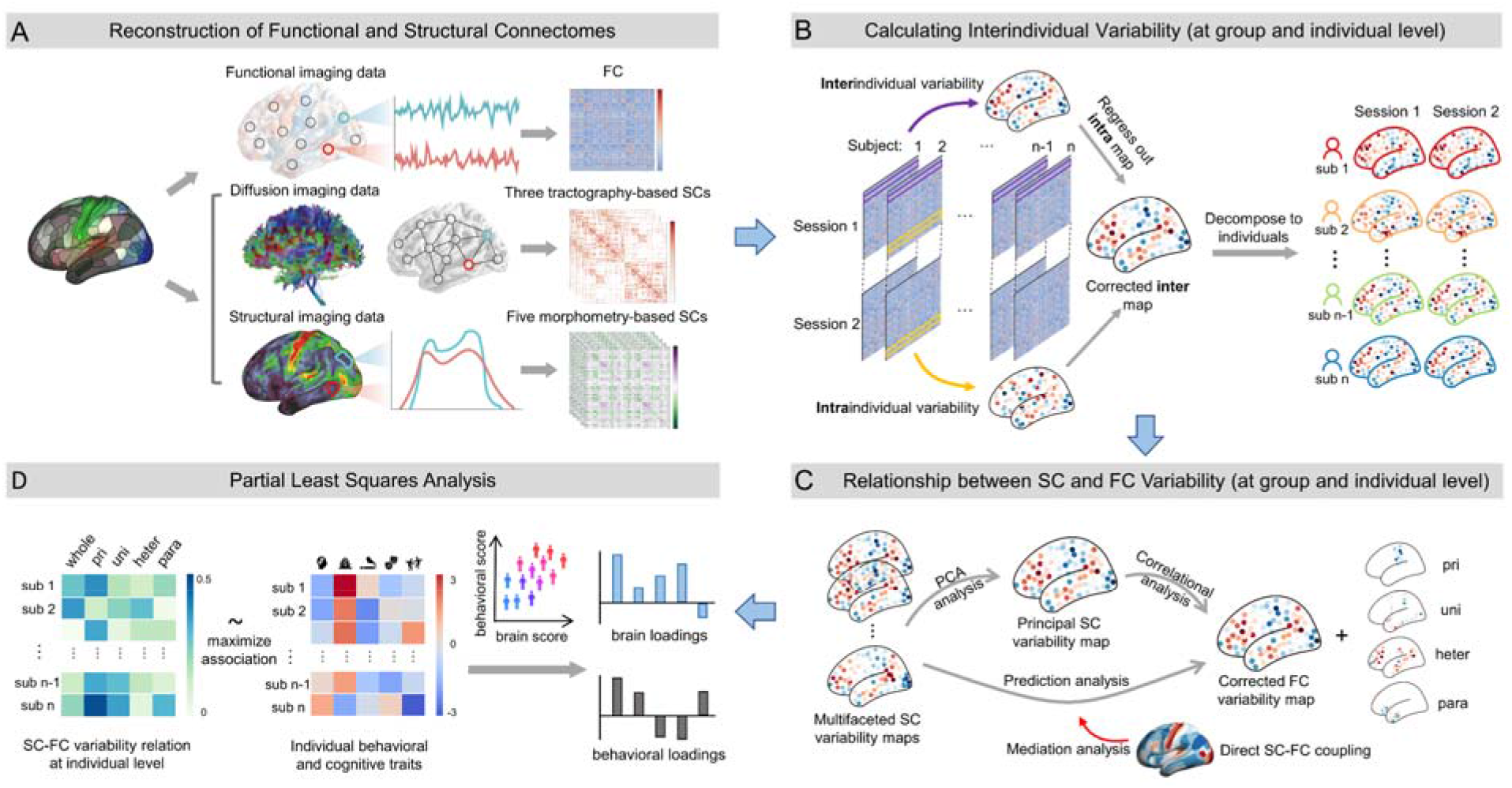
Study overview. (A) Functional connectome (FC) and structural connectome (SC) reconstruction. FCs were obtained by computing Pearson’s correlation coefficients among the time series of all pairs of nodes. Tractography-based SCs and morphometry-based SCs were obtained by computing internode probabilistic white matter fiber streamlines and internode gray matter morphometric similarity, respectively. (B) Following the approach proposed by Mueller and colleagues (Mueller et al., 2013), we calculated the adjusted interindividual FC and SC variability and decomposed the group-level variability pattern into individual unique contributions. (C) Both linear and nonlinear computational analyses were used to test the hypothesis that FC variability is structurally constrained across the whole brain and each hierarchical system. We hypothesize that direct SC-FC coupling may underlie the alignment between SC and FC variability. pri, primary cortex; uni, unimodal cortex; heter, heteromodal cortex; para, paralimbic cortex. (D) Partial least-squares (PLS) analysis was used to explore the multivariate correlations between FC-SC variability correspondence at the individual level and multiple cognitive and behavioral traits.

For intraindividual variability, we observed that the spatial patterns were nonuniformly distributed throughout the cortical mantle, in which both FC and SC variability values were prominent primarily in the medial temporal lobe and insular cortex (Fig 2A). However, we observed significant differences in the intraindividual variability values among these connectomes (one-way repeated analysis of variance (ANOVA), F = 2916.6, p < 0.0001): the highest variability values in FC, followed by morphometry-based SC, and the lowest variability values in tractography-based SC (post hoc pairwise comparisons analysis, all p < 0.0001, Bonferroni-corrected).

**Figure 2.**
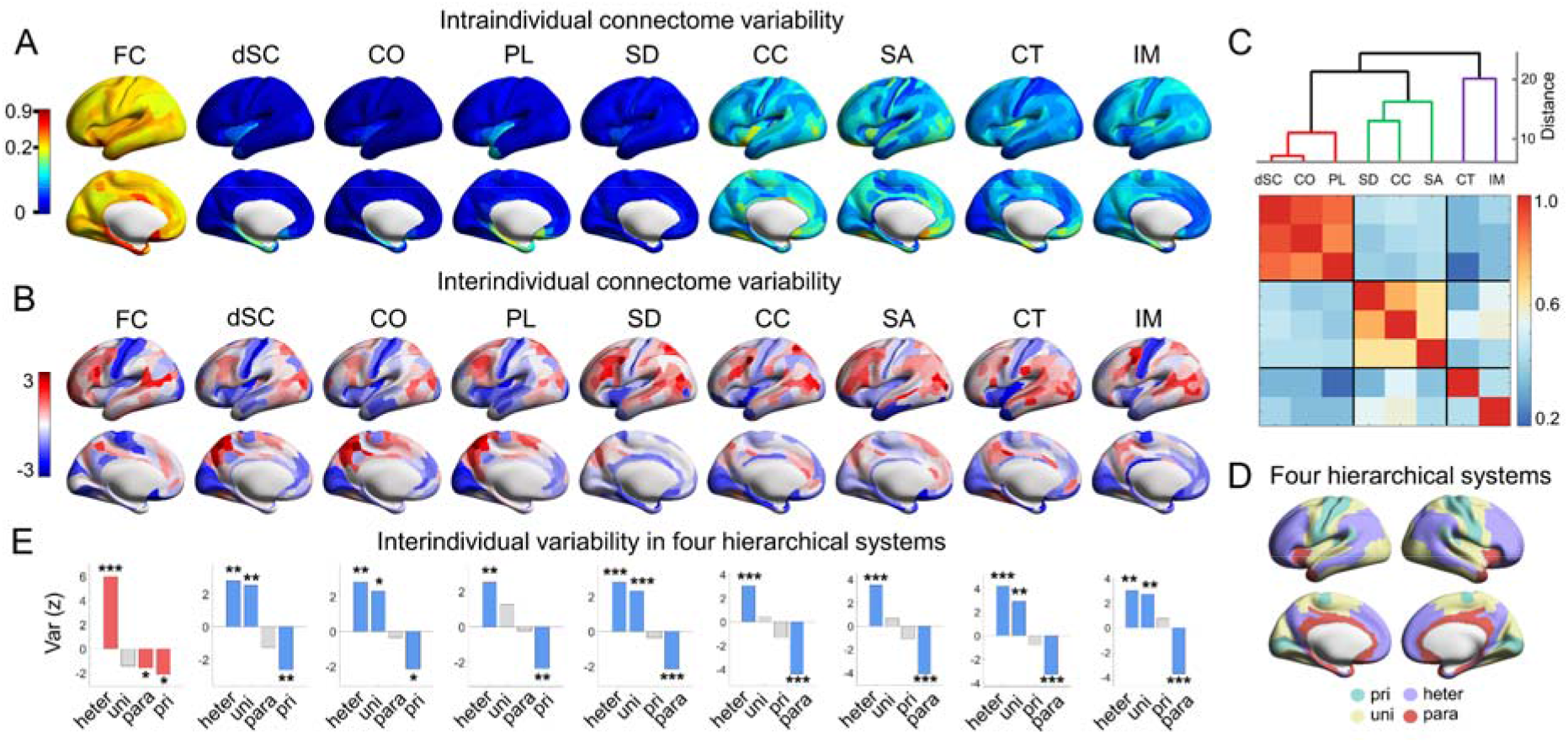
The heterogeneous spatial distribution of FC variability and SC variability. (A) Intraindividual FC variability and SC variability of all anatomical features. The results showed here are for the left hemisphere, and the relatively symmetrical whole-brain patterns are displayed in Fig S1A. (B) Interindividual FC and SC variability after accounting for intraindividual variability. The raw variability values were scaled using a rank-based inverse Gaussian transformation (Van der Waerden, 1952) for visualization. Negative scores represent raw values below the average. The interindividual FC and SC variability pattern without regressing intraindividual variance is shown in Fig S1B. (C) Results of hierarchical clustering analysis and the feature-feature correlational matrix. (D) The cortical hierarchy assignments for each region in Glasser’s 360-atlas (primary, green; unimodal, yellow; heteromodal, purple; paralimbic, red) (Liu et al., 2020; Mesulam, 1998). (E) Interindividual variability across four hierarchical systems. Nodewise variability values are averaged according to their hierarchical classes. To determine these variabilities were not driven by spatial autocorrelation, we performed a spatial permutation test by spinning class positions 10,000 times as a null model. The class-specific mean variability values were expressed as z scores relative to this null model. A positive z score indicated greater variability value than expected by chance. Classes with significant z scores are shown in color, and those with nonsignificant z scores are shown in gray. FC, functional connectome; dSC, direct structural connectome; PL, path length-based SC; CO, communicability-based SC; SD, sulcal depth-based SC; CC, cortical curvature-based SC; SA, surface area-based SC; CT, cortical thickness-based SC; IM, intracortical myelination-based SC. * p < 0.05; ** p < 0.01; *** p < 0.001. Values of a brain map were visualized on the inflated cortical 32K surface (Glasser et al., 2016) using BrainNet Viewer (Xia et al., 2013).

For interindividual variability, FC variability values (Fig 2B, first column) were higher in the lateral prefrontal cortex and temporal-parietal junction and lower in sensorimotor and visual regions, which is in line with previous studies (Mueller et al., 2013; Gao et al., 2014; Xu et al., 2018; Stoecklein et al., 2019). Interindividual SC variability patterns were generally similar to those in FC variability patterns, but there were some feature-specific distributions in several regions, such as the precuneus cortex, lateral prefrontal cortex, and temporal-parietal junction cortex (Fig 2B, second to last column). Hierarchical clustering analysis divided these SC variabilities into three clusters (Fig 2C), representing distinct structural signatures in tractography- and morphometry-based (cortical folding-based and cortical architecture-based, separately) connectomes. To further test whether spatial distributions of FC and SC variabilities follow a hierarchical manner from primary to heteromodal organization, we first classified all 360 brain nodes into four cortical hierarchies (Mesulam, 1998; Liu et al., 2020) (Fig 2D) and calculated the mean interindividual variability within each class. Then, we used a spherical projection null test by permuting class positions 10,000 times to estimate whether these variabilities were determined by the hierarchical classes or derived by spatial autocorrelation (Alexander-Bloch et al., 2018; Liu et al., 2020). We found that the heteromodal class displayed significantly greater mean variability than expected by chance (p _spin_ < 0.0001 for FC; p _spin_ < 0.01 for all SCs; Fig 2E). In contrast, the primary class (p _spin_ < 0.05 for FC; p _spin_ < 0.05 for all tractography-based SCs) and paralimbic class (p _spin_ < 0.05 for FC; p _spin_ < 0.001 for all morphometry-based SCs) displayed significantly lower variability than null models.

### 3.2. The Structural Constraints of the Hierarchical Organization of Functional Variability

Next, we sought to determine the relationship between individual FC and SC variability patterns using both linear and nonlinear approaches.

First, considering the distinct structural patterns provided by tractography- and morphometry-based connectomes, we generated a linear representation of the structural variability by applying a principal component analysis to the interindividual SC variability maps that included all eight SC characteristics. The first principal component explained 54.9% of the variance in total variabilities across brain nodes (Fig 3A, left), with contributions from tractography-based SC variability, cortical folding-based SC variability, and cortical architecture-based SC variability in a descending order (Fig 3A, left inset). Similar to single-feature findings, the first principal component of structural variations was significantly higher in heteromodal and unimodal areas and lower in primary and paralimbic areas than expected by chance (all p _spin_ < 0.05, Fig 3A, right). Next, we calculated the spatial correlation between functional variability and principal structural variability at both the whole-brain and system levels. We found a significant spatial correlation across whole-brain nodes (adjusted r = 0.43, p _spin_ < 0.0001, Fig 3B, left). At the system level, the spatial correlations were greatest in primary regions (adjusted r = 0.71, p _spin_ < 0.001) and smallest in heteromodal regions (adjusted r = 0.30, p _spin_ < 0.05) (Fig 3B, right).

**Figure 3.**
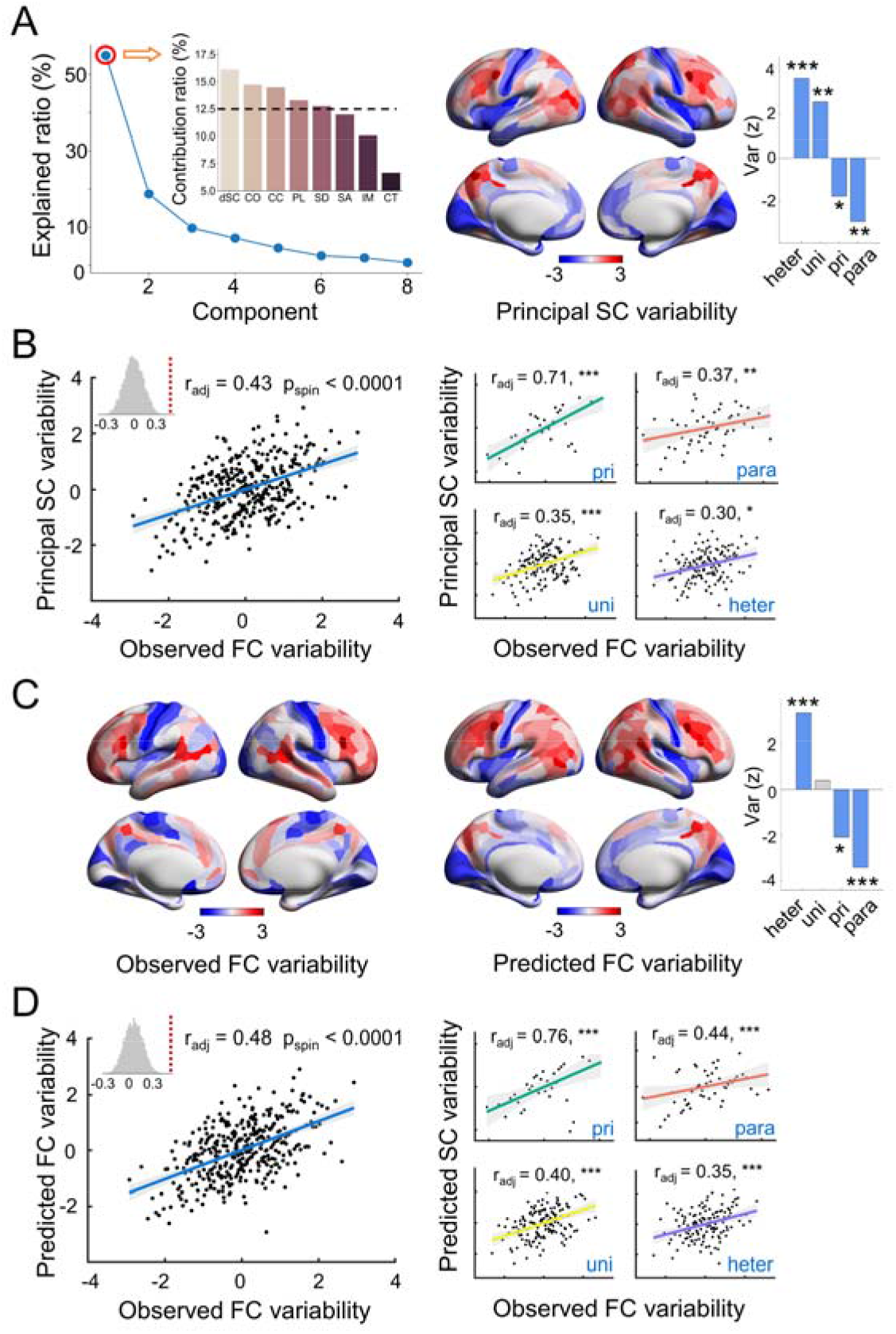
The relationship between interindividual FC and SC variability at the population level. (A) Principal component analysis (PCA) estimated linear combinations of SC variability patterns with maximum variance across the cortical mantle. The first component accounted for 54.9% of the total variance, with the largest weight contribution of tractography-based SC variability, followed by cortical folding-based SC variability and cortical architecture-based SC variability (Inset Figure) (left panel). The dashed line on the inset figure indicates the expected average contribution (1/number (features) = 12.5%). A feature with a contribution percent larger than this cutoff could be considered important in contributing to the principal component. The principal SC variability varied across cortical regions, exhibiting significantly higher values in heteromodal and unimodal areas but significantly lower values in primary and paralimbic areas (p _spin_ < 0.05) (right panel). (B) The correlational relationships between principle SC variability and FC variability were significant in the whole brain (left panel) and each hierarchical system (right panel) with highest values in the primary system and lowest values in the hetermodal system. (C) The observed FC variability map and the predicted FC variability map that was obtained by a nonlinear SVR model. (D) The predicted FC variability was also significantly associated with the observed FC variability in the whole brain (left panel) and each hierarchical system (right panel) with highest values in the primary system and lowest values in the hetermodal system. * p < 0.05; ** p < 0.01; *** p < 0.001.

Second, to further test whether the FC variability architecture can be predicted from unifying SC variability patterns, we performed a multivariate nonlinear-kernel-based prediction analysis using a supervised support vector regression (SVR) model. This model took eight SC variability maps as input and was trained and estimated in a leave-one-out strategy. We found that FC variability (Fig 3C) could be significantly predicted across whole-brain nodes (adjusted r = 0.48, p _spin_ < 0.0001, Fig 3D, left). At the system level, we found that the predictive power of SC variability for FC variability decreased along the hierarchy axis (adjusted r = 0.76, p _spin_ < 0.0001 in the primary cortex; adjusted r = 0.40, p _spin_ < 0.001 in the unimodal cortex; adjusted r = 0.44, p _spin_ < 0.01 in the paralimbic cortex; adjusted r = 0.35, p _spin_ < 0.001 in the heteromodal cortex, Fig 3D, right). Taken together, these findings indicate that interindividual FC variability was structurally constrained by SC variability in a primary-to-heteromodal hierarchical order.

### 3.3. Structure-Function Coupling Mediates the Relationship between Individual Structural and Functional Variability

The connection patterns of brain FCs are formed by interactions of neuronal elements via complex structural pathways (Wang et al., 2015b). Direct SC-FC coupling is treated as a basic index for the intensity of structural constraints on brain function and has been considered to reflect common cortical hierarchical organization (Suárez et al., 2020). Let us assume that for each individual, if the SC profiles of brain nodes were exactly coupled with their FC profiles (with Pearson’s coefficient = 1), the topography of individual FC and SC variability patterns would be the same. Thus, it is reasonable to hypothesize that this SC-FC coupling may underlie the alignment between interindividual SC and FC variability. To test this hypothesis, we first computed the direct SC-FC coupling for each given node using a multilinear regression approach (Vazquez-Rodriguez et al., 2019). Consistent with previous findings (Baum et al., 2019; Vazquez-Rodriguez et al., 2019, Zamani Esfahlani et al., 2022), we found that the sensorimotor and occipital cortices exhibited relatively high SC-FC coupling, while the lateral parietal, frontoparietal and temporal cortices exhibited relatively low SC-FC coupling (Fig 4A). Using a bootstrapped mediation analysis, we found that the spatial pattern of direct SC-FC coupling partially mediated the relationship between principal structural variability and functional variability (indirect effect *ab*: β = 0.05, p < 0.01, 95% confidence interval = [0.02, 0.09], bootstrapped n = 10000; Fig 4B). The spatial pattern of direct SC-FC coupling was significantly negatively associated with both functional variability and principal structural variability (path *a*: β = −0.16, p = 0.002; path *b*: β = −0.32, p < 0.001), indicating that brain nodes with stronger SC-FC coupling correspond to those with weaker interindividual variability. Consistent results were also found when using the predicted functional variability map driven by SVR analysis instead of the principal structural variability map (Fig 4C). These results suggest that the constraint of interindividual structural variability on functional variability was mediated by direct SC-FC coupling.

**Figure 4.**
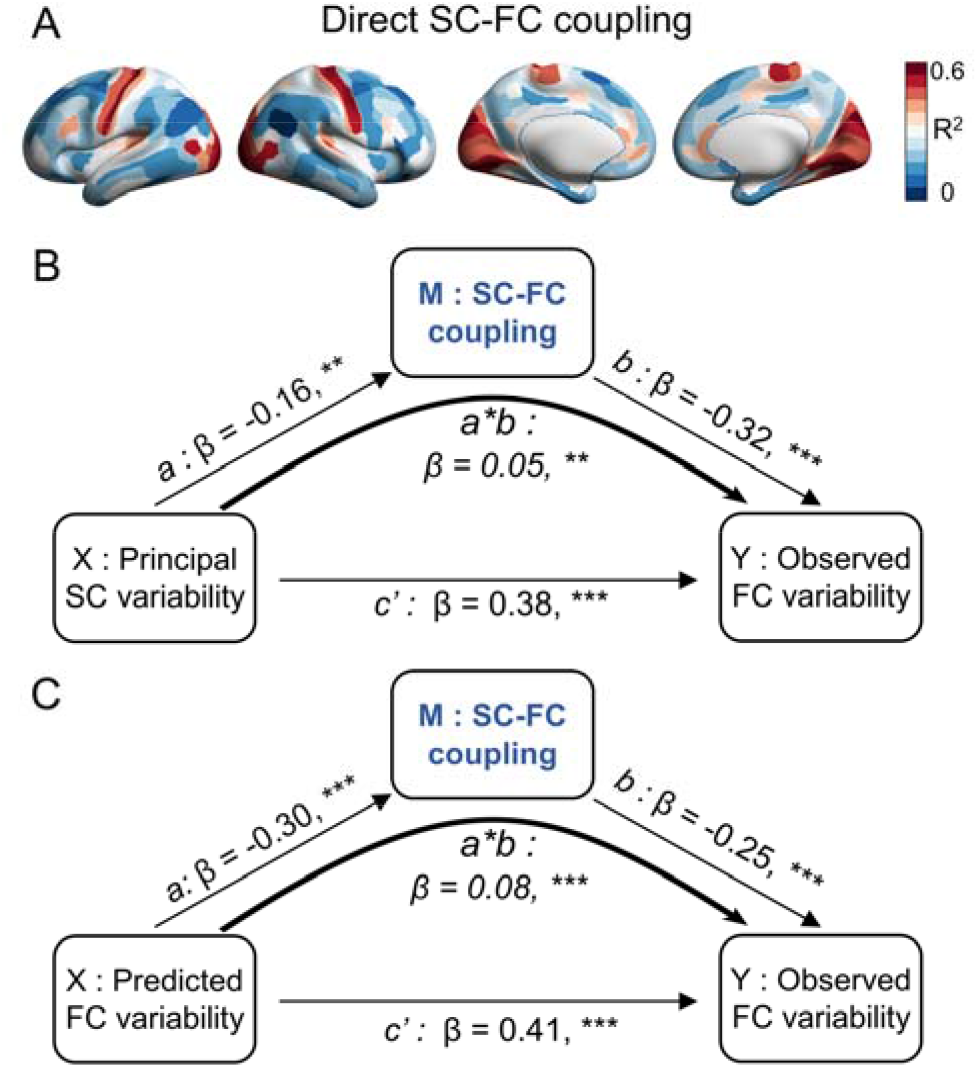
The SC-FC coupling pattern mediates the relationship between SC and FC variability. (A) Nodal differences in SC-FC coupling. For each node, the coupling of structural and functional connectivity profiles was estimated by a multilinear regression framework (Vazquez-Rodriguez et al., 2019). (B) SC-FC coupling partially mediated the correspondence between principal SC variability and observed FC variability. Path *a*: β = −0.16, **, CI: [−0.26, −0.06]; path *b*: β = −0.32, ***, CI: [−0.38, −0.24]; path *c*’: β = 0.38, ***, CI: [0.29, 0.49]; path *a*b*: β = 0.05, **, CI: [0.02, 0.09]. (C) SC-FC coupling partially mediated the correspondence between predicted FC variability and observed FC variability. Path *a*: β = −0.30, ***, CI: [−0.41, −0.20]; path *b*: β = −0.25, ***, CI: [−0.32, −0.17]; path *c*’: β = 0.41, ***, CI: [0.32, 0.52]; path *a*b*: β = 0.08, ***, CI: [0.05, 0.11]. The significance of the mediation effect was identified using 95% bootstrapped confidence intervals (bootstrapped n = 10,000). ** p < 0.01; *** p < 0.001.

### 3.4. The Robust Correspondence between Individual Functional and Structural Uniqueness

The group-level FC or SC variability pattern captures the total variance within a particular population, while it is unable to represent the unique contributions of each individual. To characterize the individualized source of group-level variability, we decomposed the overall interindividual FC or SC variability map into individual uniqueness patterns. Briefly, the individual uniqueness delineates the personal deviation of individual connectivity profiles from the populational connectivity profiles while controlling for intraindividual variance (detailed in Materials and Methods). Similar to the group-level analysis, for each individual, we performed principal component analysis on all structural uniqueness patterns to represent the individual principal structural uniqueness map. The intraindividual spatial similarity of either functional unique maps or principal structural unique maps (functional uniqueness: r, mean ± standard deviation (std) = 0.80 ± 0.09; principal structural uniqueness: r, mean ± std = 0.91 ± 0.02) were significantly higher (ps < 0.0001) than interindividual spatial similarity from either session (functional uniqueness: r, mean ± std = 0.52 ± 0.05; principal structural uniqueness: r, mean ± std = 0.67 ± 0.02) (Fig 5A, 5B). Consistent results at the system level are shown in Fig S2. An individual identification analysis further revealed that the success rates of functional uniqueness and principal structural uniqueness were 91.7% and 100%, respectively. In addition, to be expected, the interindividual standard deviation for both functional and structural uniqueness was smallest in the primary sensorimotor and visual regions and largest in the heteromodal cortex (Fig 5C). These results indicated that the individual uniqueness maps of the brain could act as potential signatures to describe the intersubject connectome diversity.

**Figure 5.**
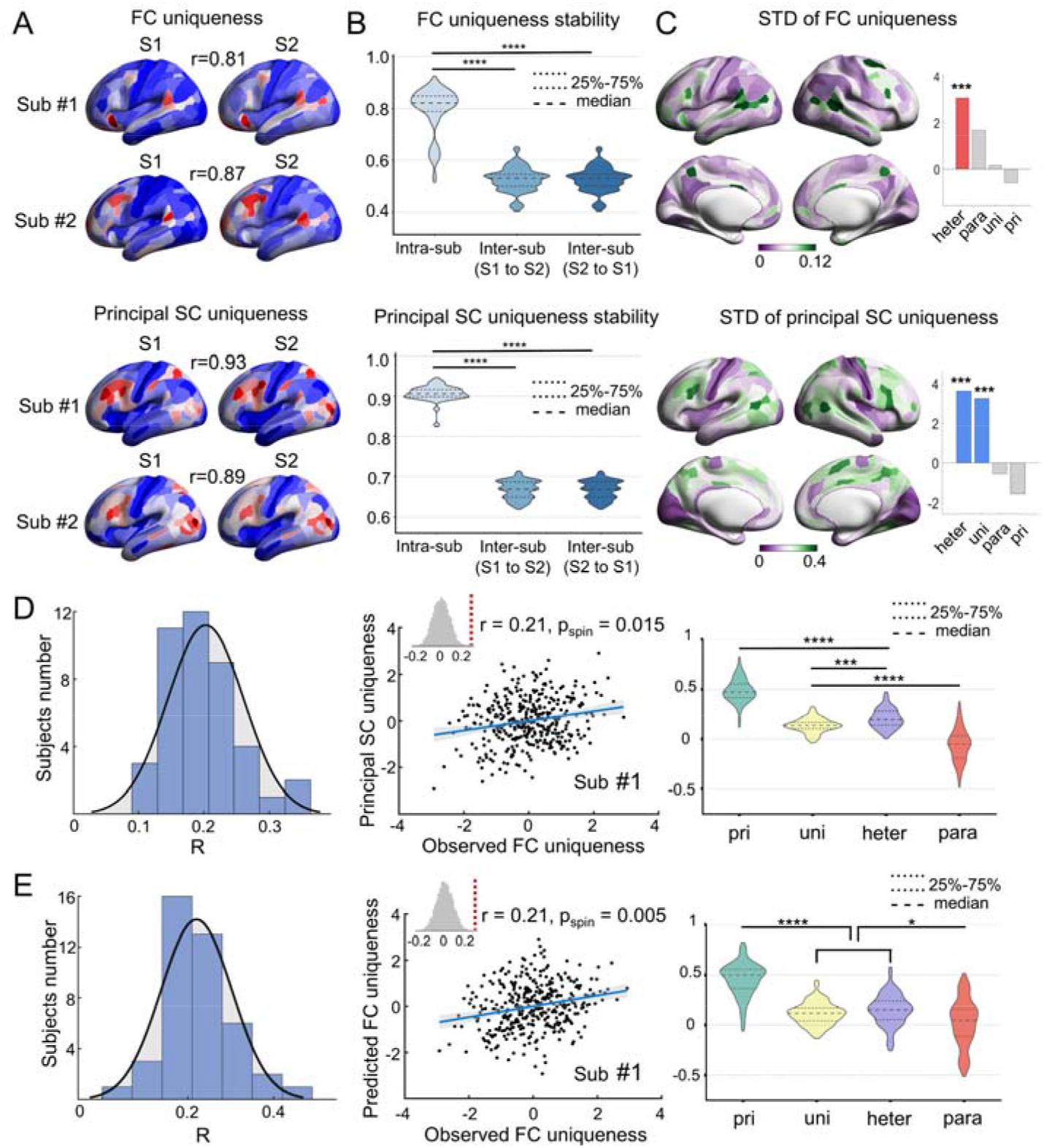
Within-individual FC and principal SC uniqueness patterns and hierarchical correspondence between individual FC uniqueness and SC uniqueness. (A) The FC uniqueness (top panel) and principal SC uniqueness pattern (bottom panel) of two example subjects (#1 and #2) showed high reliability across sessions. (B) The whole brain spatial similarity within the same subject (intrasub) of either FC uniqueness maps (top panel) or principal SC uniqueness maps (bottom panel) was significantly higher than those between different subjects (intersub, Session 1 to Session 2 and Session 2 to Session 1 were shown separately). The spatial similarity between two uniqueness maps was estimated using Pearson’s correlation coefficient. The significances of differences between categories were tested using a nonparametric permutation test. (C) The standard deviation (std) of FC and principal SC uniqueness map across 42 subjects were shown in left. Hetermodal system showed significant higher std than expected by chance (right). (D) The spatial similarities between FC uniqueness and principal SC uniqueness at whole-brain (left panel, distribution of 42 subjects; middle panel, an example of subject #1) and system level (right panel). The spatial similarities were higher in the primary system than the hetermodal system: one-way repeated analysis of variance (ANOVA), F = 190.88, p < 0.0001, post hoc pairwise comparisons analysis, all ps < 0.001. (E) The consistent results estimated by the nonlinear SVR model. *, p < 0.05; ***, p < 0.001; ****, p < 0.0001.

We next focused on the spatial relationships between the functional and structural uniqueness maps using correlational analysis and predictive models as previously described. We found that the spatial correlations between functional uniqueness and principal structural uniqueness across the whole brain were moderate (r ∈ [0.09, 0.36], mean r = 0.20; Fig 5D, left) but significant for most subjects (p _spin_ < 0.05 for 39 of 42 subjects, other 3 subjects were marginally significant (p _spin_ ∈ [0.05,0.09])).

The significant correlation of an example subject was shown in a scatter plot (Fig 5D middle). At the system level, the spatial correlations were largest in the primary cortex, followed by the heteromodal, unimodal cortices and paralimbic cortex (Fig 5D, right). These spatial relationships were also confirmed by the nonlinear SVR model (Fig 5E). Together, these results show that the structure-function correspondence of individual uniqueness was robust across subjects.

### 3.5. The Structure-Function Correspondence of Brain Uniqueness Reflects Individual Cognitive and Behavioral Traits

Studies have shown that the extent of direct SC-FC coupling at the individual level is associated with cognitive performance, such as cognitive flexibility (Medaglia et al., 2018) and executive function (Baum et al., 2019). The alignment between SC and FC uniqueness reflects the degree of the individual-specific constraint rule out of group constraints and could also support individual cognitive traits. To address this issue, we performed a partial least-square (PLS) analysis to evaluate the latent relationship between the structural-functional alignments of brain connectome uniqueness and general behavioral traits across individuals. For brain measurements, we considered the structural-functional linear correspondences at both the whole-brain and system levels. For behavioral domains, we used individual behavioral measurements in five independent dimensions derived from 109 behavioral items in the HCP S1200 dataset (Tian et al., 2020). After performing PLS analysis, we decomposed the brain-behavior associations into driving latent components (LCs), which are the optimal linear combinations of original brain or behavioral measurements. We found that the first LC (LC1, accounting for 62.6% of the covariance) significantly survived after permutation tests (p = 0.004), with a significant association (r = 0.11, p < 0.01) between brain and behavioral composite scores (Fig 6A). We further obtained the brain and behavioral loadings of this component by calculating Pearson’s correlations between the original measurements and their composite scores. A large absolute loading value indicates great importance. We found significant positive brain loadings in structural-functional alignments, with the greatest loading value at the whole-brain level and ascending loading values from unimodal to heteromodal systems (Fig 6B). The behavioral significant loadings included tobacco use, the personality-emotion score, and the mental health score (Fig 6C). We observed consistent results using the nonlinear SVR model (Fig S3). These results suggest that the hierarchical correspondence between functional and structural uniqueness, especially in the heteromodal cortex, reflects the individual traits in human general behaviors.

**Figure 6.**
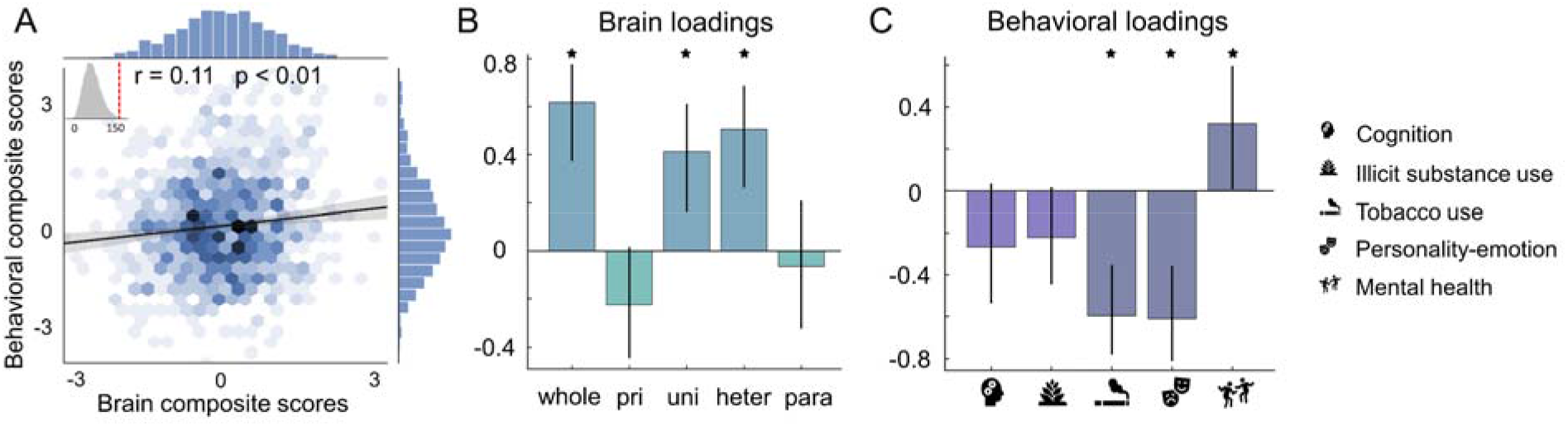
Partial least-square analysis revealed a multivariate association between the structural-functional alignment of individual uniqueness and individual behavioral traits. (A) Correlation between brain and behavioral composite scores of 954 subjects. The inset figure shows the null distribution of singular values estimated by the permutation test (n=10,000). The dashed red line represents the actual singular value estimated for the first latent component. (B) The brain loadings that were calculated by the correlations between subjects’ brain measurements and their brain composite scores. (C) The behavioral loadings that were calculated by the correlations between subjects’ behavioral measurements and their behavioral composite scores. Error bars indicate bootstrapping fifth to 95th percentiles, and robust results are indicated by a star symbol. The consistent results based on the nonlinear SVR model are shown in Fig S3.

### 3.6. Sensitivity and Replication Analyses

To estimate the effect of the sample size (Button et al., 2013; Dubois and Adolphs, 2016) of the HCP TRT dataset, which contains only 42 subjects, we validated all analyses using a large-sample dataset of HCP S1200 release (1012 subjects). Because this dataset contains repeat-scan fMRI data but no repeat-scan diffusion or structural imaging data, we calculated the individual structural variability without accounting for intrasubject variability. This approach was acceptable since we found that the principal structural variability with removing intrasubject variability was highly similar to that without removing intrasubject variability (r = 0.94, p _spin_ < 0.0001, Fig S4) using the HCP TRT dataset. However, the correlation value between functional variability with the removal of intrasubject variability and that without the removal of intrasubject variability was only 0.41 (p _spin_ < 0.001, Fig S4), indicating the importance of excluding intrasubject variations when estimating intersubject FC variability. Over the large sample, we observed highly consistent results for the hierarchical relationships between brain structural and functional variability, the mediation effect of direct structure-function coupling, and the significant correspondence at the single-subject level (Figs S5-S8).

Next, we estimated the reliability of our results by considering several confounding factors. First, the number of nodes in the Glasser360 atlas assigned to each of the four hierarchical systems is not equal (range: 32-144). To verify that the differential correspondence of variability maps between hierarchical systems is not influenced by the node number, we randomly selected brain regions of a fixed number of 30 in all systems 1000 times and recomputed the correlation coefficient based on these subsets of nodes (Fig S9). Second, to reduce the partition effect, we consistently validated the results in another well-known cortical system—the seven cytoarchitectonic classes described by Von Economo and Koskinas (Von Economo and Koskinas, 1925) (Fig S10). Third, to rule out the possibility that the spatial variability maps are caused by the lower fidelity of cross-subject alignment, we assessed the relationship between the spatial variability patterns and the surface deformation map that occurred in cross-subject registration. We found that the subject-averaged surface deformation map was not associated with functional variability (r = −0.04, p _spin_ > 0.05) but significantly associated with principal structural variability (r = −0.18, p _spin_ < 0.05). After regressing out the cross-subject registration deformation map, the functional variability pattern and principal structural variability pattern were still significantly correlated (r = 0.43, p < 0.0001; Fig S11). Fourth, a recent study excluded one of all twins in HCP dataset while estimating a brain-behavior association (Zekelman et al., 2022). To exclude the influence of the related subjects, we excluded 17 subjects (one of the twins) from the TRT dataset and found that the individual variability pattern of both FC and SC were almost unchanged (r > 0.97 for all variability patterns). Meanwhile, we observed highly consistent results for the correspondence between structural and functional variability at the whole-brain level and system level (Fig S12).

Although several processing steps are used to remove the motion effect in the current work (including ICA-FIX denoising, motion parameter regression, and motion scrubbing), it’s still valuable to estimate the influence of subjects with high-level motion. To this end, we excluded subjects with high-level head motion for either fMRI or dMRI images in both HCP TRT dataset and S1200 dataset and further validated all main results. Specifically, the head motion indexes (i.e. mean and mean absolute deviation of the frame-to-frame displacements) of each subject during the fMRI and DWI scanning sessions were estimated (by using Movement_RelativeRMS.txt from minimal fMRI preprocessing pipeline and using eddy_unwarped_images. eddy_movement_rms from minimal dMRI preprocessing pipeline). After that, we excluded subjects with one or more indexes greater than 1.5 times the inter-quartile range of the corresponding index distribution (Zamani Esfahlani et al., 2022). Under this criterion, five subjects were excluded for the HCP TRT dataset, while 95 subjects were excluded for the HCP S1200 dataset. Validation analysis was carried out by using the remaining 37 subjects (TRT dataset) and 917 subjects (S1200 dataset). Of note, the individual variability patterns of both FC and SC were almost unchanged before and after excluding subjects with high-level head movements (r > 0.98 for all variability patterns). The correspondence between SC and FC variability at the whole-brain level and system level and the brain-behavior multivariate association were highly repeatable as well (Figs S13 and S14).

## 4. Discussion

In this study, we demonstrate for the first time that FC individual variability is largely constrained by unifying SC variability and that this constraint follows a hierarchical pattern with stronger coupling in the primary cortex and weaker coupling in the heteromodal cortex. By decomposing group-level brain variability patterns into individual uniqueness maps, we also demonstrate individual-level structural-functional uniqueness correspondences that act as indicators to capture individual cognitive traits. These results markedly advance our understanding of the structural underpinnings behind individual variability in both FC and cognitive performance.

The SCs and FCs of the human brain are inherently correlated. Despite extensive research on SC-FC coupling (Suárez et al., 2020), whether interindividual FC variability is anatomically constrained by SC variability is still an open question. We emphasize that the characterization of this relationship depends on several key factors. First, a relatively large sample size with repeated structural and functional brain scans is needed. Previously, Camberland et al. (Chamberland et al., 2017) and Karahan et al. (Karahan et al., 2021) both reported no significant correlation between SC and FC variability at a global level, which could be attributable to the small sample sizes of their studies (Chamberland et al., 2017; Karahan et al., 2021) or the confounds of intrasubject variations (Karahan et al., 2021). In this study, we found that the correlation between FC variability maps before and after regressing intraindividual variance was only 0.41 (0.94 for principal SC variability), highlighting the necessity of removing intraindividual FC variance when studying the relationship between FC and SC variability. Second, the unifying contribution from multimodal connectome signatures, including tractography- and morphometry-based networks as well as communication models, should be taken into account. Although Karahan et al. reported a significant spatial correlation (ρ = 0.32) between FC variability and tractography-based SC variability (Mansour et al., 2021), this study ignored the contribution of communication models and morphometry-based SC: the former provides important anatomical scaffolding for interregional communication through multistep signal propagation (Avena-Koenigsberger et al., 2017; Crofts and Higham, 2009) and manifests as strong interregional FC (Goni et al., 2014; Suárez et al., 2020), and the latter is sensitive for capturing axo-synaptic projections within the same cytoarchitectonic class, which is not characterized by diffusion MRI (Goulas et al., 2017; Seidlitz et al., 2018) and partly recapitulates interregional FC (Alexander-Bloch et al., 2013; Geng et al., 2017). It is widely recognized that multifaceted but integrated approaches provide complementary advantages and perspectives to explore human brain organization (Paquola et al., 2020; Van Essen et al., 2019). Using repeated-measurement, multimodal connectome features in a large sample from the HCP database, in the present study, we highlight the constraints of integrated SC variabilities on FC variability. Notably, these results are only based on correlation or prediction analyses, which could not exclude potential confounders such as parcellation. Future investigations employing neurodynamic models (Demirtas et al., 2019) might be helpful to reveal underlying determinants.

It is worth noting that the present study exhibited a hierarchical correspondence between SC and FC variability at both the group and individual levels. These findings agree with mounting evidence that the primary-to-heteromodal hierarchy has been emphasized as a unifying principle for brain structural-functional organization (Huntenburg et al., 2018), including intracortical myelination (Huntenburg et al., 2017), cortical laminar interareal projections (Paquola et al., 2019) and SC-FC couplings (Zamani Esfahlani et al., 2022). The hierarchical correspondence between SC and FC variability might have several originations. First, from an evolutionary view, to adopt rich environmental conditions, the heteromodal cortex is untethered from canonical sensory-motor activity cascades during cortical expansion to form varied and complex wiring organizations (Buckner and Krienen, 2013), such as inherently spatially distributed communities (Margulies et al., 2016) and long-range rich-club edges (Griffa and van den Heuvel, 2018; van den Heuvel and Sporns, 2011). Simulation and neuroimaging studies demonstrated that these wiring characteristics facilitate the high complexity of functional dynamics (Senden et al., 2014; Zamora-Lopez et al., 2016), which may result in inconsistencies in the alignment between SC and FC variability. Second, the heteromodal cortex processes mixed and diverse signals from multiple sources (Avena-Koenigsberger et al., 2017; Betzel et al., 2018), which vary greatly across individuals (Betzel et al., 2018). These signals may form discrepant FC profiles across individuals under similar structural organizations. Third, accumulating evidence has shown that the regional heterogeneity of individual variability is largely determined by both i) genetic factors, including heritability (Anderson et al., 2021; Valk et al., 2022) and gene expression (Li et al., 2021a), and ii) environmental factors, such as socioeconomic status (Leonard et al., 2019) and chronic stress (Sheth et al., 2017) during development (Foulkes and Blakemore, 2018; Tooley et al., 2021). Heteromodal regions undergo prolonged maturation compared with primary regions (Cao et al., 2017; Gilmore et al., 2018; Zhao et al., 2019), which could provide a high freedom of plasticity for functional and structural refinements during the postnatal environment and form varying FC-SC alignments across individuals. Previous works have demonstrated the heterogeneous age-related changes in regional SC-FC coupling (Baum et al., 2019; Zamani Esfahlani et al., 2022). Further exploration on whether and how SC-FC variability correspondence changes during neurodevelopment and aging is critical to understand the origination of hierarchical correspondence between brain SC and FC variability.

We observed a robust mediation effect of SC-FC coupling on the SC-FC variability correspondence, which indicated that the alignment of SC and FC variability may be intrinsically affected by the direct structure-function relationship. This finding agrees with previous evidence on the origin of brain variability. A simulation study on the SC-FC relationship has shown that the spatial distribution of FC variability could be derived by the heterogeneous oscillations of neurons through interregional SC pathways (Demirtas et al., 2019). Other studies have demonstrated that the flexible dependence of underlying SCs enables FCs to reconfigure flexibly, which provides key information for individual identification (Griffa et al., 2022) and results in interindividual cognitive switching variability (Medaglia et al., 2018). Other biological sources may also underlay the correspondence of SC and FC variability. For instance, we also observed a significant medication effect of regional cerebral blood flow (Fig S15), which has been demonstrated to be highly coupled with regional metabolism during brain development (Raichle and Mintun, 2006) and to affect the structural-functional relationship (Chen et al., 2021). How the SC-FC variability correspondence originates still requires further detailed investigation. Crucially, identifying genetic, environmental or biological influences between FC and SC variability will be important for neuroscientific studies of individualized clinical applications.

The individual brain connectome can be regarded as a common backbone with unique personal refinements (Gratton et al., 2018; Wang et al., 2021; Zimmerman et al., 2018). By decomposing the group-level variability patterns into individual deviation maps, we delineated the SC and FC variability at the individual level. These maps show high session-to-session stability and high identification accuracy of individuals, reflecting the individual uniqueness of the brain. Other efforts have also been made to extract the individual refinements by comparing individual deviations to group-level normative distributions during disorder (Liu et al., 2021; Marquand et al., 2016) and development (Nadig et al., 2021). These unique measurements show crucial significance in brain fingerprinting (Seitzman et al., 2019) and individual cognitive and behavioral predictions (Nadig et al., 2021; Saggar et al., 2015). Notably, the correspondence of individual SC and FC uniqueness also followed a hierarchical axis and showed a significant association with general behavioral and cognitive performance, with most contributing features deriving from the association cortex. This finding is in line with previous studies showing that the association cortex acts as the most informative predictor of individual fluid intelligence (Finn et al., 2015) and other cognitive and behavioral domains (Mueller et al., 2013). Recent studies have pointed out that study on brain-behavior associations requires large cohorts containing thousands of samples to obtain reproducible results (Marek et al., 2022). Although our findings are based on cohorts of near one thousand (958 subjects) and remain significant after removing subjects with high-level head motion (870 subjects), future validations on larger-sample and multi-site datasets (such as UK Biobank) are still desirable. Meanwhile, we adopted a doubly multivariate PLS analysis, in which brain systems are jointly mapped into several general behavioral traits. This approach enables the representation of interindividual variability in brain and behavior and improves reproducibility (Marek et al., 2022; Genon et al.,2022). Additionally, the hierarchical contributions of brain systems for the behavioral associations, that is the highest in heteromodal system, emphasized the need to include regional-specific effects in brain-wide association studies (Gratton et al., 2022). For detailed behavioral measurements, we found that the tobacco use shows highest loadings than other behavioral scores after PLS analysis. Prior studies also found that tobacco use is relatively highly correlated with brain measurements than mental health and illicit substance use dimensions (Mansour et al., 2021; Wang et al., 2021; Seguin et al., 2021). In addition, Illicit drugs use and cognition showed no significant brain association. Illicit drugs use is highly related to the corticostriatal functional circuits (Ersche et al., 2020). The lack of subcortical connections in the current FC-SC uniqueness evaluation may hinder the seeking for regional connectivity support of it. Total cognition score has been shown to be mostly related to the regional SC-FC coupling of only 4 focal brain regions (located in bilateral middle cingulate cortex and supplementary motor area) (Gu et al., 2021). Future estimations with fine-grained regional parcellation and nodal level indicators are needed to capture its reliable regional association. Our measurements of the correspondence between SC and FC uniqueness reflect the pure individual constraints between SC and FC that rule out group factors, which could offer insights into reflecting the true interindividual diversity of human behaviors and cognitions and highlight the potential for progress in individualized clinical therapeutics and interventions.

Several issues need to be considered. First, although the HCP TRT dataset contains data from only 42 subjects, it is the largest public young-adult database with high-quality and full repeat-scan multimodal images, which provides an irreplaceable opportunity to explore the constraints of structural variations on FC variability. In the future, we hope to replicate our findings in a larger-sample dataset. Second, it is intriguing to explore the potential genic origins of the spatial correspondence between structural and functional variability. However, this exploration would depend on the availability of an individual-specific gene expression dataset with data from a large number of donors. Third, a recent study emphasized a regional “model preference” for SC-FC relationship (Zamani Esfahlani et al., 2022). Diversified SC communication models can be adopted for different brain regions in future studies on the SC-FC relationship of individual variability. Fourth, there are inherent limitations to reconstruct accurate anatomical projections from dMRI-based tractography (Maier-Hein et al., 2017; Reveley et al., 2015; Thomas et al., 2014), such as the underestimation of long-distance white matter fiber bundles (Reveley et al., 2015) and the missing of tiny fiber tracts (Maier-Hein et al., 2017). These could bring biases to the observed SC-FC relationships of individual variability. How to reduce tractography biases in the estimation of the true regional structure-function correspondence, especially in terms of individual variability, needs further research. Finally, a recent study delineated that FC variability changes diversely in neuropsychiatric disorders (Sun et al., 2020), and future studies should focus on exploring whether the SC-FC correspondence of individual variability transforms in brain disorders, especially for individualized diagnosis and treatment evaluation.

## Acknowledgments

We thank Dr. Ye Tian for generously sharing the preprocessed behavioral data. We also thank Dr. Xi Yu for the discussion on behavioral data. This study was supported by the National Key Research and Development Project (2018YFA0701402), National Natural Science Foundation of China (82021004, 81801783, 82071998), China Postdoctoral Science Foundation (2020TQ0050), Beijing Nova Program (No Z191100001119023), and Fundamental Research Funds for the Central Universities (2020NTST29). Imaging data were provided by the Human Connectome Project, WU-Minn Consortium (Principal Investigators: David Van Essen and Kamil Ugurbil; 1U54MH091657) funded by the 16 NIH Institutes and Centers which support the NIH Blueprint for Neuroscience Research; and by the Mc-Donnell Center for Systems Neuroscience at Washington University.

## Data and Code Availability

MRI data are redeposited and publicly available in the HCP ConnectomeDB (https://db.humanconnectome.org/). Intermediate data supporting the results are available at https://github.com/sunlianglong/Structural-Insight-into-Individual-Functional-Variability.

Codes used for the neuroimaging analysis and the statistical models are available at https://github.com/sunlianglong/Structural-Insight-into-Individual-Functional-Variability.

